# *PNuts*: Global spatio-temporal database of plant nutritional properties

**DOI:** 10.1101/2022.08.29.505708

**Authors:** Fabio Berzaghi, Francois Bretagnolle, Zwanga Ratshikombo, Alya Ben Abdallah

**Affiliations:** Laboratory of Climate and Environmental Sciences (LSCE) - UMR CEA/CNRS/UVSQ; Gif-sur-Yvette, France; Sasakawa Global Ocean Institute, World Maritime University, Fiskehamnsgatan 1, 211 18 Malmö, Sweden; UMR CNRS/uB 6282 Biogéosciences, Université de Bourgogne, 6 bd Gabriel, 21000 Dijon, France

**Keywords:** nutritive values, nutritional values, chemical defenses, plant-animal interactions, ecological modelling

## Abstract

Plant nutritional properties (crude protein, fibers, minerals, and carbohydrates) and chemical defenses (tannins and phenols) are key traits determining food quality and feeding preferences of animals and humans. Plant nutritional properties are also relevant to crop production and livestock. Here we present *PNuts*, a global database containing > 1000 species and > 13,000 records of nutritional properties of different plant organs complete with location and time of collection (year/month/season). Species include crops and wild plants and are classified in six functional groups: legumes, herbs, grasses, lianas, shrubs and trees. Plant organs include leaf, fruit, seed, stem, twigs, flower, root, and bark. *PNuts* data can be used as inputs for ecological analyses and model parametrization requiring large amounts of data. *PNuts* provides an important tool to better understand the importance of nutritional properties in plant eco-physiology and the implications for humans and animals’ food quality and plant-animal interactions in a context of global changes.

## Introduction

Plants are an important direct and indirect source of nutrients for livestock, wild herbivores, and humans. Plants vary considerably in their content of protein, fiber, carbohydrates, minerals, and chemical defenses (1). Different proportions of these biomolecules, also called nutritional properties or values, determine the nutritional quality (digestibility and palatability) of plants for animals and humans (1–3). Nutritional quality is linked to reproductive success, weight gains, milk production and behavior (1). In grazing systems for example, it is essential to evaluate forage quality to know the amounts of protein and energy supplied by forages to sustain livestock and dairy production.

Nutritional properties can vary significantly among plant functional groups (1). For example, legumes contain greater quantities of proteins while grasses and trees are an important source of fibers and minerals (4, 5). Variations in nutritional contents are also observed across plant organs, as these perform specific eco-physiological functions and thus differ chemically and structurally (6). For example, leaves usually have more cell solubles (protein, sugars, and lipids) needed for photosynthesis than cell wall constituents such as stems. Additional important differences are found across development stages with young leaves having higher nutritive quality and digestibility compared to mature leaves, which are less palatable due to more investment in structural and chemical defenses (4, 7).

Nutritional traits vary not only between plant species, functional groups, organs and development stages, but also as a function of biotic and abiotic environmental conditions. Differences can be observed within same plant species collected at different locations with varying climate (8). Changes in forage nutrient contents may also be related with changes in soil nutrient contents and fluctuations of soil carbon concentrations(9).

Despite a large body of research on plant nutritional properties, there is no centralized and open-access database providing geolocated and temporal records of plant nutritional properties of different plant organs covering a wide taxonomic range in different habitats.

Here we present *PNuts,* a global database of the most studied biomolecules and traits that determine the nutritional quality of plants covering a broad range of temporal, spatial, and ecological conditions (Figs. 1 and 2). Nutritional properties in *PNuts* are reported by plant organ and species are classified in six functional groups: legumes, herbs, grasses, lianas, shrubs and trees (Figs. 2 and 3). *PNuts* aims to energize collaborations between ecology, conservation biology, environmental sciences, climate and ecological modelling, crop and livestock science. These data are required to better understand the importance of nutritional properties in plant eco-physiological processes and the implications for ecosystems, agriculture, and economy in a rapidly-changing world.

**Fig. 1.**
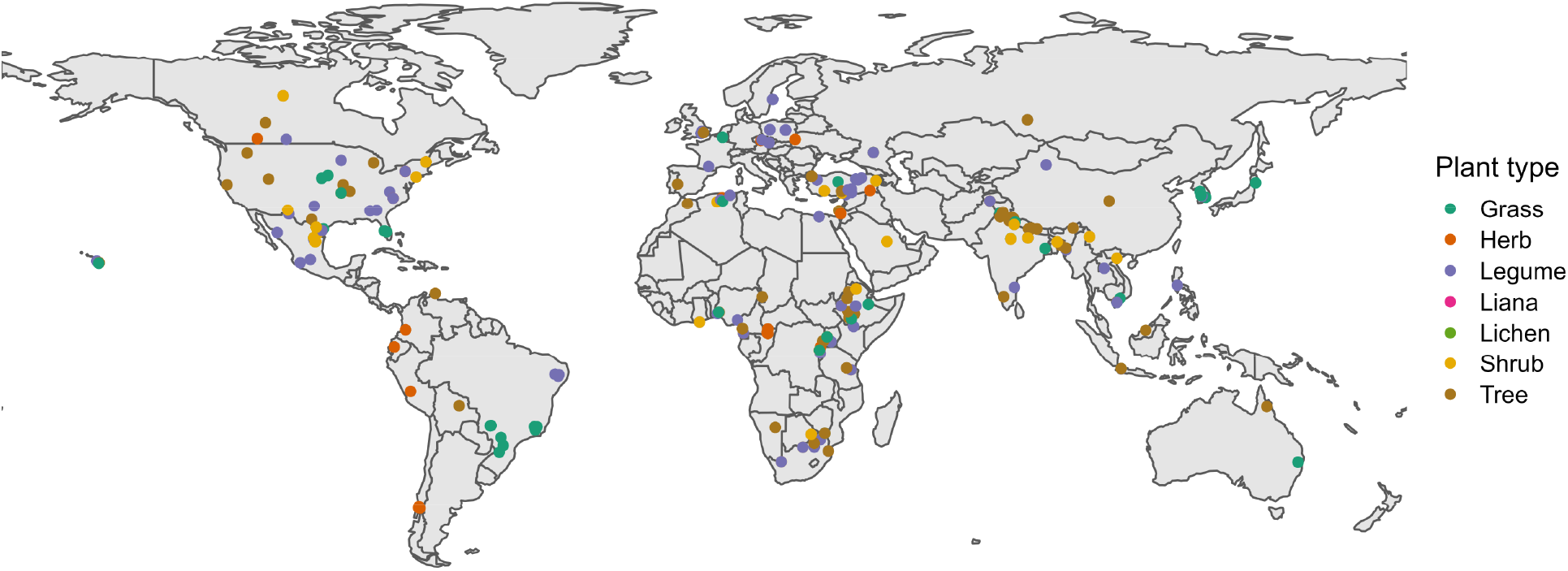
Spatial distribution of data in *PNuts* by plant functional group. See Table 1 for number of records for each functional group. Not all records in the database have a defined geographic location.

**Fig. 2.**
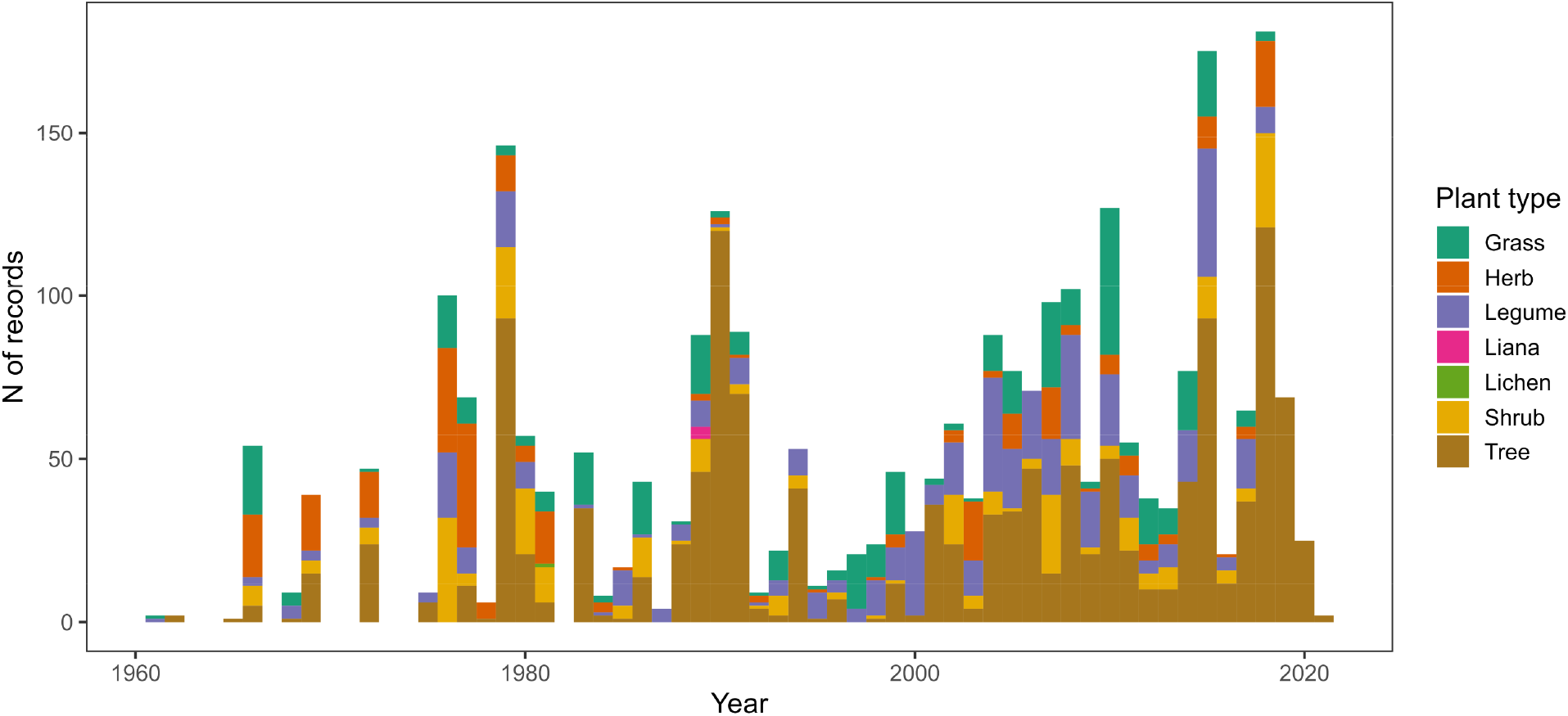
Temporal distribution of data in *PNuts* by plant functional group.

**Fig. 3.**
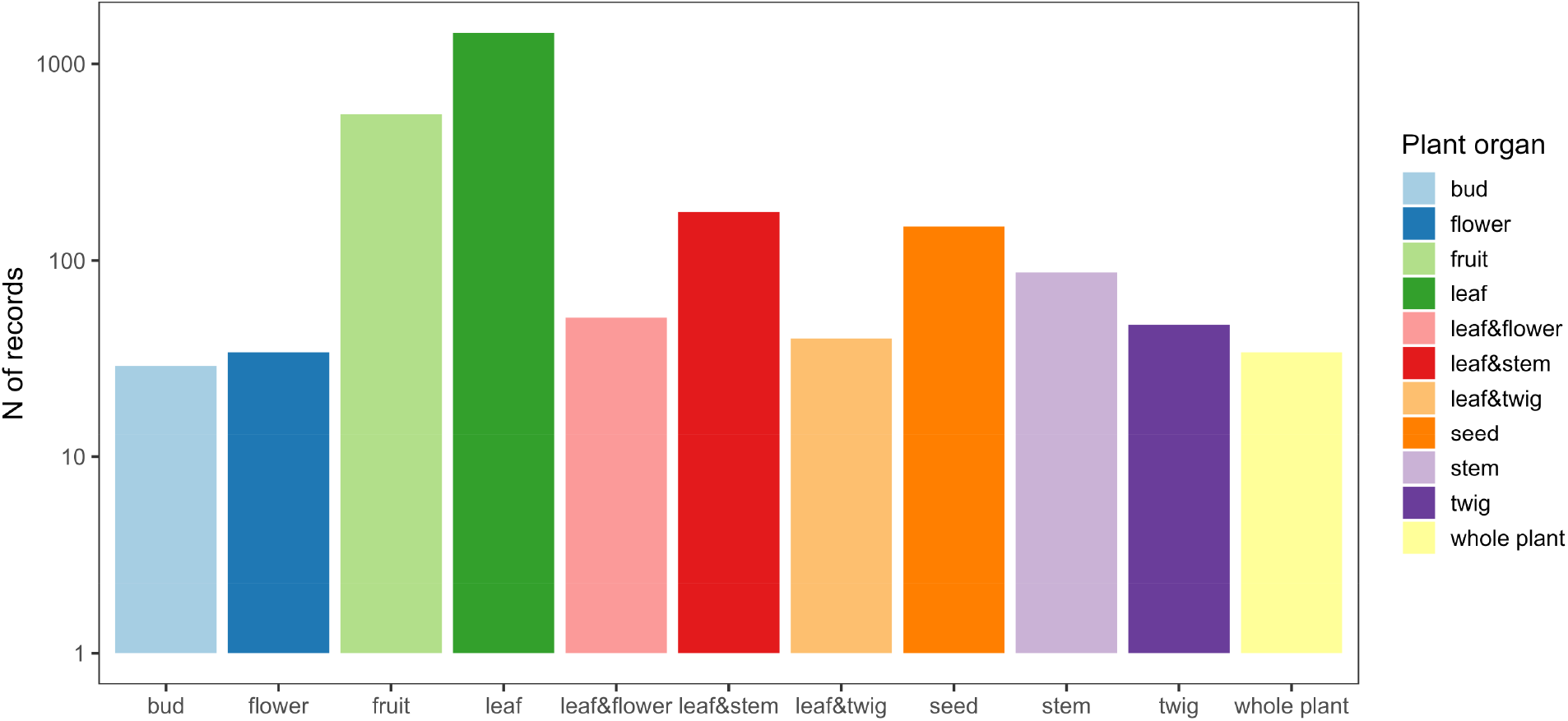
Data count of the plant organs or combination thereof with the greatest number of records. Note that the distance between the y-axis ticks is in log scale.

## Methodology

### Data collection

Data was collected primarily from peer-reviewed articles and secondarily from non-peer-reviewed publications such as PhD thesis and reports. Articles were selected in Web of Science with these keywords in a AND/OR query: (“digestibility” OR “crude protein” OR “forage quality” OR “nutritional metrics” OR “nutritive value” OR “nutritional value”) AND (“flower” OR “flowers” OR “leaf” OR “leaves” OR “twig” OR “twigs” OR “seeds” OR “seed” OR “fruit” OR “fruits” OR “roots”) AND (“herbivor*” OR “ungulate*” OR “grazer*” OR “browser*” OR “mix-feeder*” OR “livestock” OR “domestic animals” OR “captive animals” OR “pig” OR “pigs” OR “bush pig” OR “potamochoerus” OR “rodent” OR “sus scrofa” OR “wild boar” OR “swine” OR “primate*” OR “elephant*”). The terms were decided before the beginning of the search to avoid bias. Other articles were found opportunistically or from our knowledge of the literature.

**Table 1.** Nutritional properties, taxonomy, and environmental variables contained in *PNuts*.

| Nutritional property | Variable name in *PNuts* | Unit | Variable type | N of records |
| --- | --- | --- | --- | --- |
| Dry matter | DM | % or g/100g | Continuous | 971 |
| Crude protein | CP | % of dry matter | Continuous | 2561 |
| Ash (minerals) | Ash | % of dry matter | Continuous | 1711 |
| Acid detergent fiber | ADF | % of dry matter | Continuous | 1817 |
| Neutral detergent fiber | NDF | % of dry matter | Continuous | 1553 |
| Water-soluble carbohydrates | WSC | % of dry matter | Continuous | 152 |
| Total non-structural carbohydrates | TNC | % of dry matter | Continuous | 261 |
| Fat | Fat | % of dry matter | Continuous | 405 |
| Crude tannins | CT | % of dry matter | Continuous | 473 |
| Total tannins | TT | % of dry matter | Continuous | 87 |
| Total phenols | TP | % of dry matter | Continuous | 225 |
| Digestibility | (Multiple variables, see text and Table 2) | % of dry matter | Continuous | See Table 2 |
| Energy content<br>- Gross energy<br>- Metabolizable energy<br>- Digestible energy<br>- Net energy | GE<br>ME<br>DE<br>NE | MJ/kg | Continuous | 391<br>390<br>60<br>23 |
| Plant type | Plant.type | Herb<br>Grass<br>Liana<br>Shrub<br>Tree<br>Legume | Categorical | 284<br>372<br>4<br>296<br>1394<br>546 |
| Crop | crop | Yes<br>No | Categorical | 527<br>594 |
| Plant organ (Detailed category) | Plant.part | (35 categories, see text) | Categorical | See Fig. 3 |
| Plant organ (Broad category) | Plant.part.simple | (27 categories, see text) | Categorical | See Fig. 3 |
| Longitude and latitude | longitude and latitude | Decimal degr. (WGS84) | Continuous |  |
| Elevation | elevation | Meters | Continuous |  |
| Temporal | various columns | Year/month/season | Discrete | From 1961 to 2021 |

The search led to 2758 articles which where filtered based on the relevance of the data from reading titles and abstracts and looking at the data in tables and appendixes: only 1550 articles, considered relevant, were kept. We only collected data from studies that sampled plants in their natural environment. We excluded data generated from sampling plans in modified or artificial growing environment such as greenhouses, labs, and silage or when plant parts were processed or treated beyond simple desiccation. In fertilization studies we only selected data from control/natural conditions. An average was calculated when multiple data were presented for the same species from the same site, collection time, and environmental conditions. Crude protein was included in the database if it was recorded as total crude protein, however where only nitrogen content was recorded, it was multiplied by 6.25 to calculate crude protein value in accordance with the Kjeldhal procedure (7). Plant matter digestibility was recorded, along with the reported standard deviation when available. We also recorded the digestibility measurement method used: in vitro, in vivo, in situ, in sacco or nylon bag, near-infrared reflectance spectroscopy or derived using a formula based on other chemical/nutritive properties. We recorded the animal species and age class when digestibility was estimated with live animals. Because of the different methods found in the literature to determine digestibility, we merged digestibility data into one single column (DMD) to facilitate the use of the data. The DMD column contains digestibility estimated using all methods excluding the pepsin/cellulase method because we found that this method is not commonly used. For in vitro estimations of digestibility using rumen, we recorded the rumen’s species donor. When available, we recorded the methodologies used for estimating or measuring nutritional values and energy content (E.methodology variable), which we converted to MJ/kg. To preserve uniformity between all nutritive properties, only articles where nutritive metrics were expressed as percentage of dry matter or grams per kilogram of dry matter were included.

Species were classified in six functional groups (legumes, grasses, herbs, lianas, shrubs and trees) and marked as crop (including the cultivar) or wild species according to the description provided in the text. If no description was found, we determined plan type by consulting World Flora Online, a global compendium of the world’s plant species (http://www.worldfloraonline.org/).

For each nutritional value we recorded the corresponding plant organ, which included both single organs or a combination of two or three organs. At times, particularly for grass and herbs, no specific organ was reported because the whole plant was sampled. In these cases we marked the sample as “whole plant”. We found nutritional values for a total of 35 different categories of plant organs. To facilitate the use and analysis of the data, we created a column denominated “plant.organ.simple” in which we reduce the number of plant organ categories to 27 by combining mature and immature organs, and merging “stem and sheath” (four records) with “stem”, “leaf litter” (24 records) with “leaf”, “leaf petiole and stem” (26 records) with “leaf and stem”. Plant organs with the greatest number of records are shown in Fig. 3.

When reported in the source text, we recorded the geographical coordinates of where the sample was collected with a precision of up to four decimal places. If coordinates were not reported, we used OpenStreetMap to estimate latitude and longitude based on the study site information reported in the article. If coordinates were estimated, we recorded latitude and longitude with only one decimal place, which is equivalent to an error of ± 10 km. Elevation, if not indicated in the article, was determined using digital elevation data. If the collection site was missing or could not be determined, we left latitude and longitude blank. The time of collection was recorded based on the information found in the text, or alternatively based on the publication year minus one year to account for time from collection to publication.

### Description of nutritional properties in PNuts

Dry Matter (DM): the constituents of a plant excluding water, most commonly measured by weighing the sample after letting it dry in the oven overnight. The dry matter content of a plant organ is a good indicator of the amount of nutrients being supplied to consumers (10).

All nutritional values in *PNuts* are expressed as percentage of dry matter (%DM), which in literature is also reported as the number of grams of a specific property for each 100 grams of plant dry matter. Table 1 provides an overview of all the nutritional properties, taxonomic and spatiotemporal data.

Crude Protein (CP): a crucial nutrient provided by plants to consumers as a source of energy and to perform regulatory and tissue-building functions (10).

Mineral ash (Ash): Total inorganic mineral content remaining after plant biomass is incinerated. Ash is what remains in the absence of energy, protein, and nutrients following fermentation in the rumen of an animal.

Acid Detergent Fiber (ADF): What remains after plant material has been treated under acidic conditions to remove starch, protein, fats pectin, soluble minerals, hemicellulose, oils, and free sugars. It is composed of lignin, cellulose and silica (1).

Neutral Detergent Fiber (NDF): Similar to ADF but in addition to lignin, cellulose, and silica, NDF also contains hemicellulose. Both ADF and NDF are important factors that determine plant matter digestibility (7). Fiber concentrations are considered in ration formulation because they give an indication of the quantity of forage that the animal can consume (1). A higher fiber content decreases digestibility and fiber content increases with plant growth.

Water soluble carbohydrates (WSC): Primarily composed of glucose, fructose, sucrose, and fructans, carbohydrates are a source of energy and help regulate metabolism.

Total non-structural carbohydrates (TNC): Composed of WSC plus starch.

Fat: animals require fat to perform several functions including energy storage, insulation, control metabolic function, and signaling.

Condensed Tannins (CT), Total Tannins (TT), and Total Phenols (TP): plant secondary metabolites are chemical compounds that reduce palatability and digestibility of plant biomass (10).

Plant matter digestibility: total of digestible protein, carbohydrates, cellulose, lipids, and fiber constituents of plant matter. Digestibility is mostly reported as a percentage of dry matter food intake. It can be determined from several procedures involving digestion trials with in vivo (in animal) techniques such as “in sacco” or “nylon bag”, in vitro which uses rumen liquor or pepsin/cellulase to degrade plant matter (11) (Table 2). In vitro and in vivo techniques measure the disappearance of dry matter over a period of time. Digestibility can also be estimated using equations from chemical data (11). Organic matter digestibility refers to digestibility of dry matter without the inorganic part (Ash).

**Table 2.** The different techniques used to measure plant matter digestibility and their variable name in *PNuts*. When available the standard deviation was recorded (SD). See text for brief description of the different techniques.

| Digestibility technique | Variable name in *PNuts* | Unit | Variable type | N of records |
| --- | --- | --- | --- | --- |
| In vivo (nylon bag) | NBDMD | % of dry matter | Continuous | 138 |
| Organic matter digestibility | OMD | % of dry organic matter | Continuous | 289 |
| In vitro dry matter digestibility | IVDMD | % of dry matter | Continuous | 715 |
| Pepsin/cellulase digestibility | CDIG | % of dry matter | Continuous | 221 |
| Dry matter digestibility | DMD | % of dry matter | Continuous | 1014 |
| Dry matter digestibility standard deviation | SD | % of dry matter | Continuous | 51 |

Energy content: Gross Energy (GE) is often the first step in estimating plant energy content. This measurement is often based on the heat produced from the combustion of plant matter in a bomb calorimeter (12). Digestible Energy (DE), Metabolizable Energy (ME), and Net Energy (NE) are often estimated from GE or from food nutritional values. Digestible energy is equal to GE minus energy lost in feces (12). Subtracting energy lost in urine and during digestion (gaseous emissions) from DE provides ME. Net energy is obtained by subtracting the energy lost during digestion from ME.

## Funding

This work was supported by European Union’s Horizon 2020 research and innovation program under the Marie Sklodowska-Curie grant #845265, by the French government allocation d’aide au retour à l’emploi program (FB), and by the French government aid managed by the ANR under the “Investissements d’avenir” programme with the reference ANR-16-CONV-0003 part of CLand Convengerce Institute (ZR), and by Sorbonne University Sorbonne Center for Artificial Intelligence Master’s program (ABA).

### Usage notes and data availability

*PNuts* data are currently being used for various publications which are under peer review. The data will be released gradually as the corresponding articles will be published. The dataset will become freely accessible in a spreadsheet format, which will also contain metadata and a complete list of references. An R package will also become available.

